# Identification of Novel Extracellular Vesicles Scaffold Proteins for Versatile Cargo Engineering

**DOI:** 10.64898/2026.01.04.697584

**Authors:** Tao Qiu, Rui Hu, Yuan Yi, Wenqiang Lu, Chuang Cui, Shuiqin Niu, Ke Xu

## Abstract

Extracellular vesicles (EVs) are promising drug delivery platforms that have been engineered to carry various drug modalities. Key strategies for generating those therapeutic EVs involve the direct fusion of protein of interest (POI) to EVs scaffold proteins with inherently high EVs-sorting ability. In this work, we identified raftlin (RFTN1) and poliovirus receptor (PVR) as novel EVs scaffold proteins for loading EVs lumen and surface cargos respectively. Truncation studies revealed that the N-terminus 15 residues from RFTN1 (RFTN1-N15) were sufficient for EVs engineering, as demonstrated by distinct cargos including gene editing tools, cytosolic enzymes as well as type II transmembrane proteins. On the other hand, PVR efficiently displayed secreted proteins including antibodies and serum albumins on EVs surface. Critically, RFTN1 and PVR-engineered EVs demonstrated consistent and efficient cargo delivery *in vivo*. In summary, the discovery of RFTN1 and PVR can potentially benefit EVs engineering for both fundamental research and clinical translation in the future.

## 1. INTRODUCTION

Extracellular vesicles (EVs) are naturally derived carriers that can be secreted by most cell types (van Niel, D’Angelo, & Raposo, 2018). Whereas EVs mediate cellular communications in physiological context, EVs are also undergoing intensive investigations for developing into novel therapeutic platform (Du et al., 2023; Kumar et al., 2024). Their innate properties—including low immunogenicity, ability to cross physiological barriers, and potential targeting capabilities—present attractive advantages for delivering various drug modalities (Du et al., 2023; Kumar et al., 2024).

Depending on the purposes and properties of drug modalities, the molecules of interest can be loaded either on EVs surface, or within EVs lumen (Ma et al., 2025; Yang, Xue, Duan, Mao, & Wan, 2024). Typically, EVs producer cells are genetically engineered to overexpress the protein of interest (POI) fused to an EVs scaffold protein, thereby allowing EVs cargo enrichment (Ma et al., 2025; Yang et al., 2024). Therefore, the EVs scaffold proteins with high intrinsic EVs-sorting ability are the key elements in EVs engineering strategy.

Traditionally, EVs marker proteins such as CD9, CD63 and CD81 have been applied as scaffold proteins (Ma et al., 2025; Yang et al., 2024). Over the years, several studies have performed EVs proteomic analysis to identify novel EVs scaffold proteins with higher efficiency, leading to the identification of PTGFRN and BASP1 (Dooley et al., 2021), TSPAN2 and TSPAN3 (Zheng et al., 2023) and PLXNA1 (Zhao et al., 2024). Herein, we performed proteomic studies on HEK293-derived EVs independently, and reported raftlin (RFTN1) and poliovirus receptor (PVR) as novel EVs scaffold proteins for loading EVs lumen and surface cargos respectively. Distinct cargos modalities including gene editing tools, cytosolic enzymes, type II transmembrane proteins and antibodies were tested for EVs loading and validated with desired activity both *in vitro* and *in vivo*. Therefore, we believe this study will potentially benefit EVs engineering field.

## 2. MATERIALS AND METHODS

### 2.1 Cell culture and transfection

Suspension-adapted HEK293 cell line (A23109, Quacell) was cultured in OPM-CD05 medium (81075-001, OPM Biosciences) and maintained on orbital shaker at 90 RPM in a humidified incubator at 37℃ with 8% CO2. MDA-MB-231 (CL-0150, Procell), SKOV3 (CL-0215, Procell) and B16F10 (CL-0319, Procell) were cultured in DMEM medium (11965092, Gibco) supplemented with 10% FBS (A5669701, Gibco) and maintained in a humidified incubator at 37 °C with 5% CO₂. Cells were passaged every 2–3 days.

### 2.2 EVs purification from cell culture medium

The cell culture supernatant was first filtered through 0.45 μm filter units (SLHPR33RB, Millipore), followed by centrifugation (SW32Ti, Beckman Coulter) at 133,900 g at 4 ℃ for 60 min. The crude EVs pellet were resuspended in PBS, and further layered onto 17.5% Optiprep /45% Optiprep gradient (D1556, Sigma), followed by centrifugation at 150,000 g at 4 ℃ for 16 h. The extracellular vesicles appeared as a white layer between PBS/17.5% iodixanol, which were carefully pipetted out and washed with PBS by centrifugation at 135,000 g at 4 ℃ for 3 h. The refined EVs were finally resuspended in PBS (10010, Gibco), and stored in −80℃ freezer.

### 2.3 Western blot analysis

The cells or EVs samples were first lysed with RIPA lysis buffer (R0010, Solarbio) on ice for 20 minutes. Then the total protein were quantified with MicroBCA protein assay kit (23235, Thermo scientific). After that, SDS-PAGE protein loading buffer (BL502A, Beyotime) was added into the samples and incubated at 95 °C for 10 min. Equal quantity of proteins for each sample was loaded onto 4–12% SurePAGE™, Bis-Tris gels (M00653, GenScript). After electrophoresis (170 V, 35 min), the proteins were transferred onto PVDF membrane (ISEQ00010, Millipore). The membranes were blocked with QucikBlockTM Western blocking buffer (P0252, Beyotime) for 1h at room temperature (RT) before incubation with primary antibodies overnight at 4 °C. After extensive washing with TBST wash buffer (ST673, Beyotime), the membrane was further incubated for 2h at RT with HRP conjugated secondary antibodies. Following extensive washes with TBST buffer, the membranes were incubated with Pierce ECL Western Blotting Substrate (32209, Thermo scientific) and visualized with Tannon 5200 imager (Tannon).

The following antibodies were used: CD9 (abcam, AB263019), CD81 (Cell Signaling Technology, 56039S), Calnexin (abcam, ab22595), CD63 (Cell Signaling Technology, 2897), Cre recombinase (Cell Signaling Technology, 15036), TSG101 (abcam, ab125011), TMPRSS2 (Cell Signaling Technology, 39665), 4-1BBL (Cell Signaling Technology, 59127), Arginase1 (Cell Signaling Technology, 93668), iNOS (abcam, ab178945), HRP Goat Anti-Rabbit IgG (H+L) (ABclonal, AS014), HRP Goat Anti-Mouse IgG (H+L) (ABclonal, AS003).

### 2.4 Electron Microscopy

For transmission electron microscopy, EVs were applied onto the glow-discharged copper grid (200 mesh, coated with carbon film). To perform negative staining, 2% uranyl acetate were incubated with EVs at room temperature for 1 min, followed by quick wash with distilled water to remove excess stain. The grids were air-dried before being imaged under Tecnai G2 transmission electron microscope (Thermo FEI, 120 kV).

For cryo-electron microscopy, EVs were applied to glow-discharged copper grid (Cu300, Quantifoil, 212601), and cryo-frozen by liquid nitrogen with Vitrobot plunge freezer (bolt time 4s, bolt force 0, wait time 30s). Images were acquired under cryo-EM (Thermo Glacios, 200 kV).

### 2.5 Nanoparticle tracking analysis (NTA)

To evaluate the size distribution and concentration, EVs samples were freshly diluted at 1,000–50,000 fold in 0.22-mm filtered PBS and analyzed immediately with ZetaView Nano Particle Tracking Analyzer (ParticleMetrix, PMX120-Z).

### 2.6 RT-qPCR

Total RNA from the cells plated on 96 well plates were first extracted and purified using the RNAprep Pure Micro Kit (DP420, TIANGEN). The concentration and quality of isolated RNA were detected using a NanoDrop (ThermoFisher). Reverse transcription was performed with HiScript Ill cDNA Synthesis Kit (R312-01, Vazyme). Next, the cDNA was added into the 2xSYBR green qPCR mix (A0012-R2, Ezbioscience), and quantitative PCR were analyzed with QuantStudio3 (Applied Biosystems).

### 2.7 Macrophage polarization assay

In polarization experiments, RAW264.7 cells were seeded at 20000 cells per well in 96-well flat-bottom tissue culture plates and cultured in polarizing medium overnight. M1 polarization of RAW264.7 cells was induced by 100 ng/mL Lipopolysaccharides (tlrl-3pelps, Invivogen) and 2.5 ng/mL IFN-γ (575302, Biolegend) for 24 hours. M2 polarization of RAW264.7 cells was induced by 20 ng/mL IL-4 (574302, Biolegend) for 24 hours.

### 2.8 B16F10 tumor xenograft model

Female, 8 weeks old C57BL6/J mice (GemPharmat) were implanted with 1E6 B16F10 cells/mice under the right fat pad region. When the average tumor volume reached around 60 mm^3^, the mice were randomly grouped for different treatment conditions. Intratumoral injections and tumor volume measurement were performed every day for seven days consecutively. On last day, mice were sacrificed and tumors were excised out and imaged.

### 2.9 LPS induced acute lung injury model

To establish acute lung injury model, 8 weeks old C57BL6/J mice (GemPharmat) were nasal instilled with 50 μL LPS (tlrl-eklps, InvivoGen) at 5 mg/kg dose. For EVs treatment, EVs were adminstrated by pulmonary nebulization using Micro Sprayer Aerosolizer (Y655650918, YuYanbio), at 50 μL volume in PBS per mouse. At the end of the experiment, mice were euthanized, and lung, serum and bronchoalveolar lavage fluid (BALF) samples were collected.

### 2.10 Cytokines analysis in BALF samples

The BALF supernatants were collected, centrifuged at 1000 × g for 10 min at 4°C, and cytokine levels were measured by mouse cytometric bead array (CBA) Kit (BD Biosciences). Briefly, 50 μL of samples (BALF supernatants) or known concentrations of standard samples (0–5000 pg/mL) were added to a mixture of 50 μL each of capture antibody bead reagent and phycoerythrin (PE)-conjugated detection antibody. The mixture was then incubated for 2 h at room temperature in the dark and then washed to remove unbound detection antibody. Data were acquired using a FACSCelesta cytometer and analyzed using CBA software FACP V3.0.

### 2.11 Histological analysis

Tissue samples obtained from the C57BL6/J mice were first fixed in 4% paraformaldehyde at 4 ℃ overnight, then dehydrated in 30% (w/v) sucrose solution (A610498, Sangon Biotech) for 2 days. The dehydrated samples were then embedded in Tissue-Tek O.C.T Compound (4583, SAKURA) blocks and frozen overnight. Sections acquired at 5 μm thick sections were fixed at 4% paraformaldehyde for 10 min before staining with hematoxylin and eosin (C0105M, Beyotime). Imaging analysis were performed under light microscope (MF43N, Mshot).

### 2.12 EVs proteomics identification

Nanoflow LC-MS/MS analysis of tryptic peptides from EVs was conducted on a quadrupole Orbitrap mass spectrometer (Q Exactive HF-X, Thermo Fisher Scientific, Bremen, Germany) coupled to an EASY nLC 1200 ultra-high pressure system (Thermo Fisher Scientific) via a nano-electrospray ion source. 500 ng of peptides were loaded on a 25 cm column (150 μm inner diameter, packed using ReproSil-Pur C18-AQ 1.9-µm silica beads; peptides were separated using a gradient from 8 to 12% B in 5 min, then 12% to 30 % B in 33 min and stepped up to 40% in 7 min followed by a 15 min wash at 95% B at 600 nl per minute where solvent A was 0.1% formic acid in water and solvent B was 80% ACN and 0.1% formic acid in water. The total duration of the run was 60 min. Column temperature was kept at 60 °C using an in-house-developed oven. Briefly, the mass spectrometer was operated in “top-40” data-dependent mode, collecting MS spectra in the Orbitrap mass analyzer (120,000 resolution, 350–1500 m/z range) with an automatic gain control (AGC) target of 3E6 and a maximum ion injection time of 80 ms. The most intense ions from the full scan were isolated with an isolation width of 1.6 m/z. Following higher-energy collisional dissociation (HCD) with a normalized collision energy (NCE) of 27, MS/MS spectra were collected in the Orbitrap (15,000 resolution) with an AGC target of 5E4 and a maximum ion injection time of 45 ms. Precursor dynamic exclusion was enabled with a duration of 16 s.

For data analysis, all raw files were analyzed using the Proteome Discoverer suite (version 2.4, Thermo Fisher Scientific). MS2 spectra were searched against the UniProtKB human proteome database containing both Swiss-Prot human reference protein sequences. The Sequest HT search engine was used, and parameters were specified as follows: fully tryptic specificity, maximum of two missed cleavages, minimum peptide length of 6, fixed carbamidomethylation of cysteine residues (+57.02146Da), variable modifications for oxidation of methionine residues (+15.99492Da), precursor mass tolerance of 15 ppm and a fragment mass tolerance of 0.02Da for MS2 spectra collected in the Orbitrap. Percolator was used to filter peptide spectral matches and peptides to a false discovery rate (FDR) of less than 1%. After spectral assignment, peptides were assembled into proteins and were further filtered based on the combined probabilities of their constituent peptides to a final FDR of 1%.

As default, the top matching protein or ‘master protein’ is the protein with the largest number of unique peptides and with the smallest value in the percent peptide coverage (that is, the longest protein). Only unique and razor (that is, parsimonious) peptides were considered for quantification.

### 2.13 ELISA

To determine IL15 and HSA concentration in engineered EVs, the IL15 ELISA kit (R&D systems, D1500) and HSA ELISA kit (Invitrogen, EHALB) were used according to manufacturer’s protocol. EVs were first permeabilized by incubating with lysis buffer (PBS+0.3% TritonX-100) at room temperature for 30 min, before proceeding to ELISA quantification.

### 2.14 Dye labeling of EVs and *in vivo* animal imaging

For fluorescent dye labeling of EVs, the lipophilic tracers DiO (Invitrogen, D275) and DiR (Invitrogen, D12731) were prepared according to manufacturers’ protocol. Briefly, EVs were incubated with dye (1:1000 dilution from stock) at room temperature for 30 min, and were further washed twice with PBS by centrifuge at 135,000 g for 1 h.

To analyze the distribution of EVs, the mice were I.V administered with dye-labeled EVs in 100 μl PBS through tail vein. At indicated time points, the mice were sacrificed and the internal organs (liver, spleen and tumours) were harvested and observed by an IVIS Spectrum (PerkinElmer, Waltham, MA, USA). The sampled blood was collected into the 0.5 M EDTA-treated tubes and centrifuged at 1000×g for 10 min to get the plasma for EVs detection. For EVs quantification, the plasma samples were quantified for fluorescent dye intensity by PHERAstar FSX plate reader (BMG labtech).

### 2.15 Statistical analysis

Experimental replicates were defined in the figure legends for each experiment. Statistical analyses were performed in GraphPad Prism 7 using student’s t-test for experiments with two groups or one-way analysis of variance (ANOVA) for experiments with three or more groups. Values were expressed as mean ± standard deviation (SD) or as mean ± standard error of the mean (SEM), as indicated in the figure legends. Significance labeling and p values were presented in figures descriptions.

## 3. RESULTS

### 3.1 Purification and proteomic characterization of HEK293-derived EVs

Suspension-adapted HEK293 cells grown in chemically-defined, serum-free medium were used as the EVs parental cells. We reasoned that the quality of EVs preparations critically affect the data fidelity of acquired EVs proteomics. Therefore, we first set out to develop an EVs purification process based on density gradient centrifugation. Crude EVs were refined and collected at the OptiPrep layer with density around 1.10-1.12 g/mL (Figure 1A). Nanoparticle tracking (NTA) analysis revealed EVs mean diameter to be around 120 nm (Figure 1B). The typical cup-shaped morphology was observed for EVs under transmission electron microscopy (TEM) images (Figure 1C). Additionally, minimum protein contaminants or cellular debris were observed, suggesting EVs were acquired with high-purity (Figure 1C). Western blots (WB) analysis further confirmed the purified EVs were positive for typical EVs markers CD9, CD63, CD81, TSG101 and negative for calnexin (Figure 1D).

**Figure 1.**
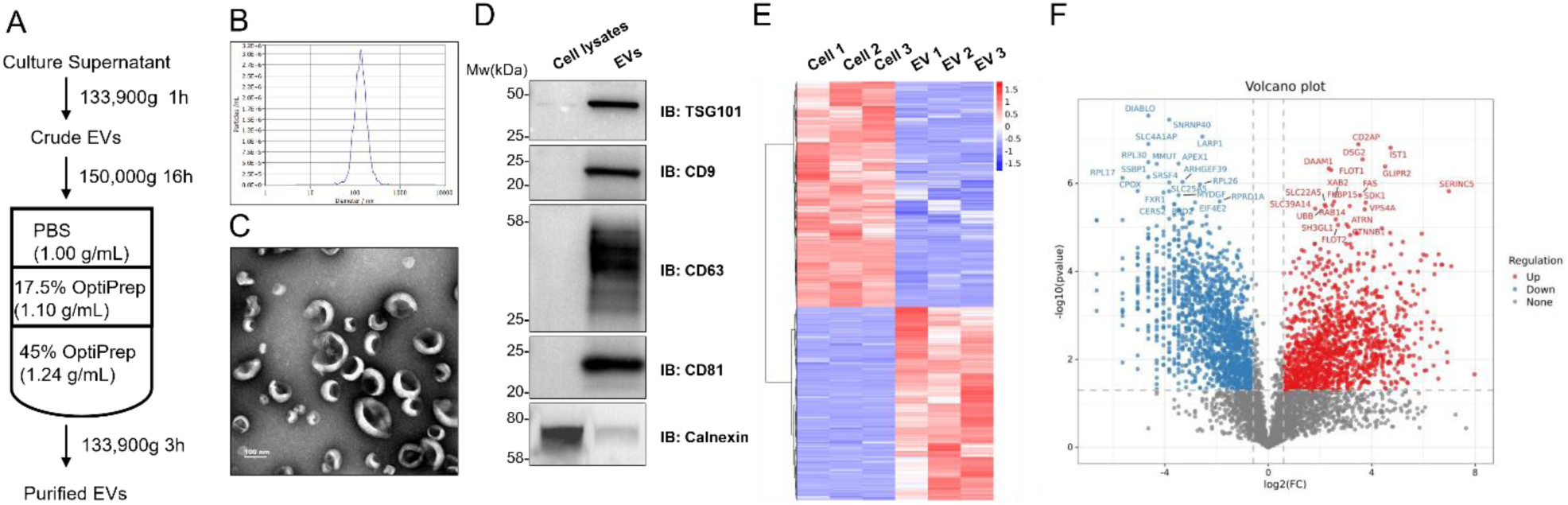
Purification and proteomic characterization of HEK293-derived EVs. (A) Schematic diagram showing the work flow for purification of HEK293-derived EVs. (B) Nanoparticle tracking analysis for purified EVs. (C) Representative transmission electron micrographs of purified EVs. Scale bar: 100 nm. (D) Western blot analysis for cell lysates and purified EVs. (E) Differential expression analysis for EVs versus cells proteomics. N=3. (F) Volcano plot showing the top candidates that were differently expressed for EVs versus cells.

Next, to identify potential EVs scaffold proteins with high EVs enrichment ability, mass spectrometry analysis was performed for purified EVs as well as parental cells. We reasoned that EVs scaffold proteins should be up-regulated in EVs proteomics versus cellular proteomics, which represented the innate EVs sorting ability. Hence, differential expression analysis was further performed for EVs versus cellular proteomics (Figure 1E-F). Protein candidates were ranked based on the significance score. Within this list, established EVs markers (CD9, CD63, CD81, TSG101) and previously reported EVs scaffold proteins (PTGFRN, BASP1, MARCKS) appeared as top candidates, confirming the good quality of proteomic data as well as the stringency of screening criteria.

Based on structural features, the top protein candidates could be further categorized as follow: (1) single-pass transmembrane proteins, including type I and type II transmembrane proteins; (2) tetraspanins; (3) multipass-transmembrane proteins; (4) membrane-associated proteins, which were anchored to membranes through lipidation modifications; (5) cytosolic proteins which typically had no association with membranes.

### 3.2 Validation of RFTN1 as EVs scaffold proteins for loading luminal cargos

We first set out to identify candidates that could be used to load cargos into EVs lumen. In previous report, BASP1 and MARCKS were favorable scaffold proteins for loading EVs luminal cargos, both proteins associated with the inner leaflet of cellular membranes through N-terminal myristoylation (Dooley et al., 2021). Similarly, the N-terminus octapeptide from Src kinases facilitated Cas9 protein encapsulation into EVs, which also depended on N-terminal myristoylation (Whitley et al., 2022). We speculated that these N-terminal lipidated proteins had unique advantages in both EVs enrichment and engineering without disturbing cargos’ function. Therefore, we focused on screening candidate proteins that had N-terminus lipidation.

RFTN1, also known as raftlin, appeared to meet the criteria among the top candidates. RFTN1 was reported as a lipid raft–associated protein that played a critical role in the organization and function of membrane microdomains essential for immune receptor signaling (Saeki, Miura, Aki, Kurosaki, & Yoshimura, 2003). RFTN1 was myristoylated on the second glycine residue, and palmitoylated on the third cysteine residue (Saeki et al., 2003). It also contained a stretch of charged residues on the N-terminus, with net charges being +2 (Figure 2A). To test the feasibility of RFTN1 to engineer cargos into EVs lumen, Cre recombinase was selected and fused to the C-terminus of RFTN1 via a flexible glycine-serine peptide linker (Figure 2B). The viral fusogen VSVG was pseudotyped on EVs membrane to facilitate Cre release (Figure 2B). A Cre-loxP recombination assay was designed such that upon successful Cre delivery, the recipient cells would emit red fluorescence in nucleus in contrast to untreated cells containing only membrane-bound green fluorescence (Figure 2C). Total 1E10 EVs particles were added to 3E4 reporter cells, followed by fluorescence analysis at 48 hours post EVs treatment. As shown in Figure 2D, RFTN1 engineered EVs successfully enriched Cre recombinase by WB analysis. Consequently, the Cre EVs delivered to recipient cells with high efficiency, recording around 70% recombination rate (Figure 2E).

**Figure 2.**
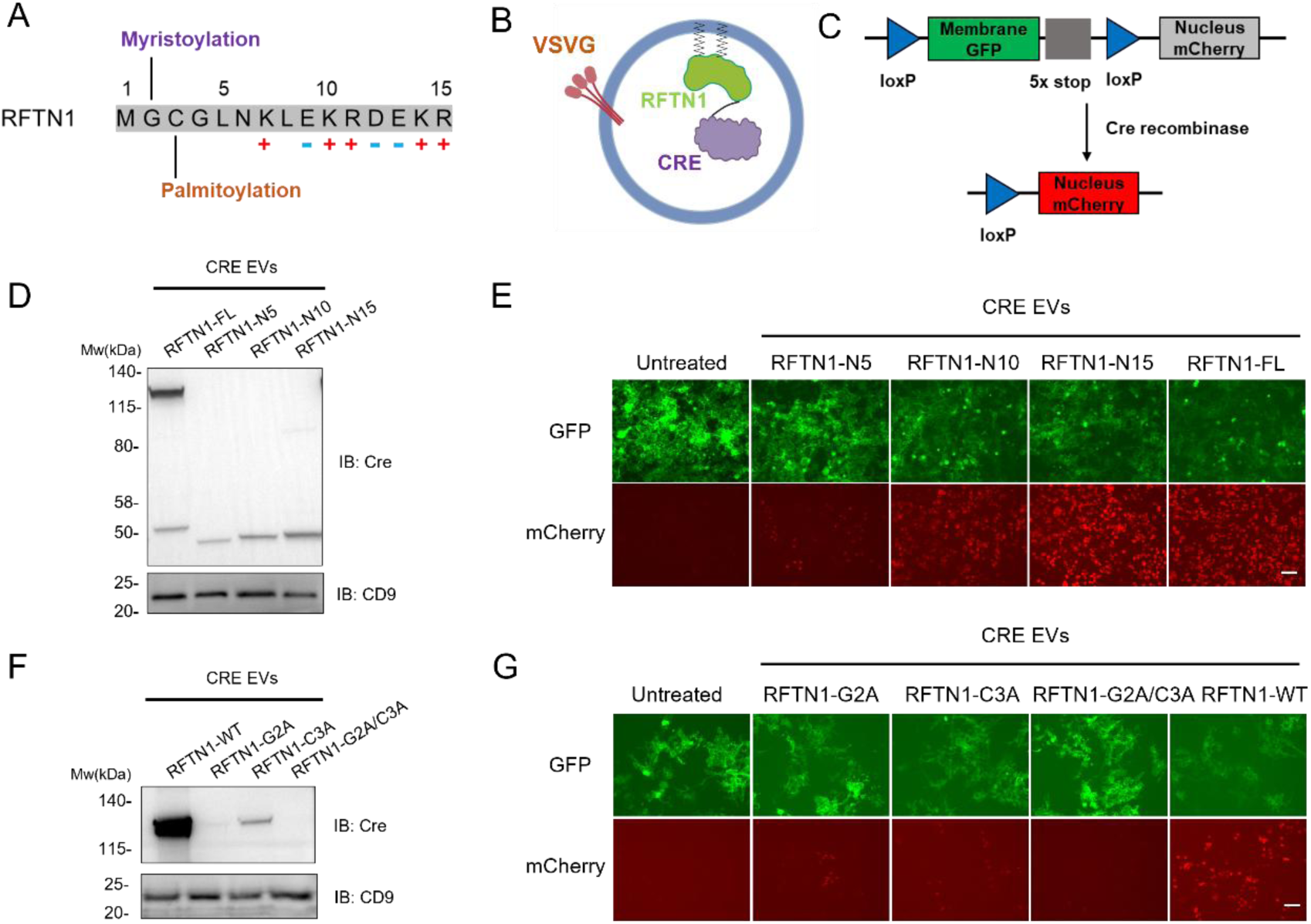
Validation of RFTN1 and derived minimum sequences as EVs scaffold proteins. (A) Sequence analysis for the N-terminus 15 amino acids from human RFTN1 protein. (B) Schematic diagram showing the EVs engineering of Cre recombinase mediated by RFTN1. (C) Schematic diagram of reporter assay designed for detecting Cre recombinase activity. (D) Western blot analysis for engineered Cre EVs from different groups, focusing on the N-terminus truncation of RFTN1. (E) Cre reporter assay analysis at 48 hours post EVs treatment. Scale bar: 20 μm. (F) Western blot analysis for engineered Cre EVs from different groups, focusing on N-terminus mutation of RFTN1. (G) Cre reporter assay analysis at 48 hours post EVs treatment. Scale bar: 20 μm.

To compare the efficiency between RFTN1 and other established scaffold proteins, BASP1 was similarly fused to Cre recombinase through GS linkers. Cre-loxP recombination assay revealed that RFTN1 performed at similar efficiency as BASP1 when added at same quantity (Supplementary Figure 1A). To test functional cargo other than Cre recombinase, a constitutive active form of β-catenin (β-catenin ΔEX3) was selected and tested (Harada et al., 1999). RFTN1 and BASP1 were fused to β-catenin ΔEX3 with GS linkers, and equal amount of EVs (1E10 particles) were added to TOPFLASH reporter cells for analysis 48 hours later. Again, RFTN1-engineered EVs efficiently activated the β-catenin signaling pathway, at similar efficiency to BASP1 as well as the small molecule lithium chloride (Supplementary Figure 1B). In summary, RFTN1 was validated as efficient scaffold proteins for engineering EVs luminal cargos.

### 3.3 Identification of the minimum sequences from RFTN1 as EVs scaffold

As reported previously, the N-terminus 10 residues from BASP1 and Src kinase were sufficient for EVs engineering and cargo loading (Dooley et al., 2021; Whitley et al., 2022). Therefore, we tested if minimum sequences could be derived from RFTN1 as well. The N-terminus 5 residues (RFTN1-N5), 10 residues (RFTN1-N10) and 15 residues (RFTN1-N15) were fused to Cre recombinase and compared for loading efficiency with RFTN1 full length (RFTN1-FL). Interestingly, RFTN1 N-terminus truncates performed gradually better as the length increased, to the point where RFTN1-N15 performed at equal efficiency as RFTN1-FL (Figure 2D-E). Although RFTN1-N5 contained the two putative N-lipidation modifications, the results suggested that the stretch of charged residues were also indispensable for EVs sorting activity.

Next, mutagenesis study was performed to confirm that the two putative N-lipidation on RFTN1 were indeed critical for its EVs sorting ability. A G2A mutation was designed to abolish the N-myristoylation, and a C3A mutation was designed to abolish the N-palmitoylation. As expected, either single mutation partially affected RFTN1’s EVs sorting activity, whereas combined double mutations almost completely abolished the activity (Figure 2F-G).

In summary, the N-terminus lipidation as well as charged residues were critical for RFTN1’s EVs sorting ability, and the N-terminus 15 aa were sufficient to be used for EVs engineering.

### 3.4 RFTN1-N15 demonstrated versatility in EVs engineering and cargo loading

Since RFTN1-N15 demonstrated sufficient efficiency in EVs engineering during proof-of-concept study using Cre-reporter system, we moved on to test on a wide range of functional cargos.

Both Cre recombinase and β-catenin ΔEX3 were nucleus-localized cargos, suggesting that RFTN1-N15’s EVs sorting signal was not mutual exclusive with cargos’ nucleus localization signal when engineered in fusion protein form. Subsequently, Cas9 protein as another typical nucleus cargo was tested for EVs encapsulation and delivery (Figure 3A). The “Stoplight” Cas9 reporter assay was designed as described previously (de Jong et al., 2020). Briefly, the successful gene editing was evidenced by EGFP fluorescence due to the correction of frame-shifted EGFP expression cassette. RFTN1-N15 was directly fused to the N-terminus of Cas9 protein, and the engineered EVs (1E10 particles) were added to 3E4 reporter cells. At 48 hours post treatment, prominent gene editing events was shown by the emerging of GFP positive cells, recording around 30% efficiency by flow cytometry quantification (Figure 3B-C).

**Figure 3.**
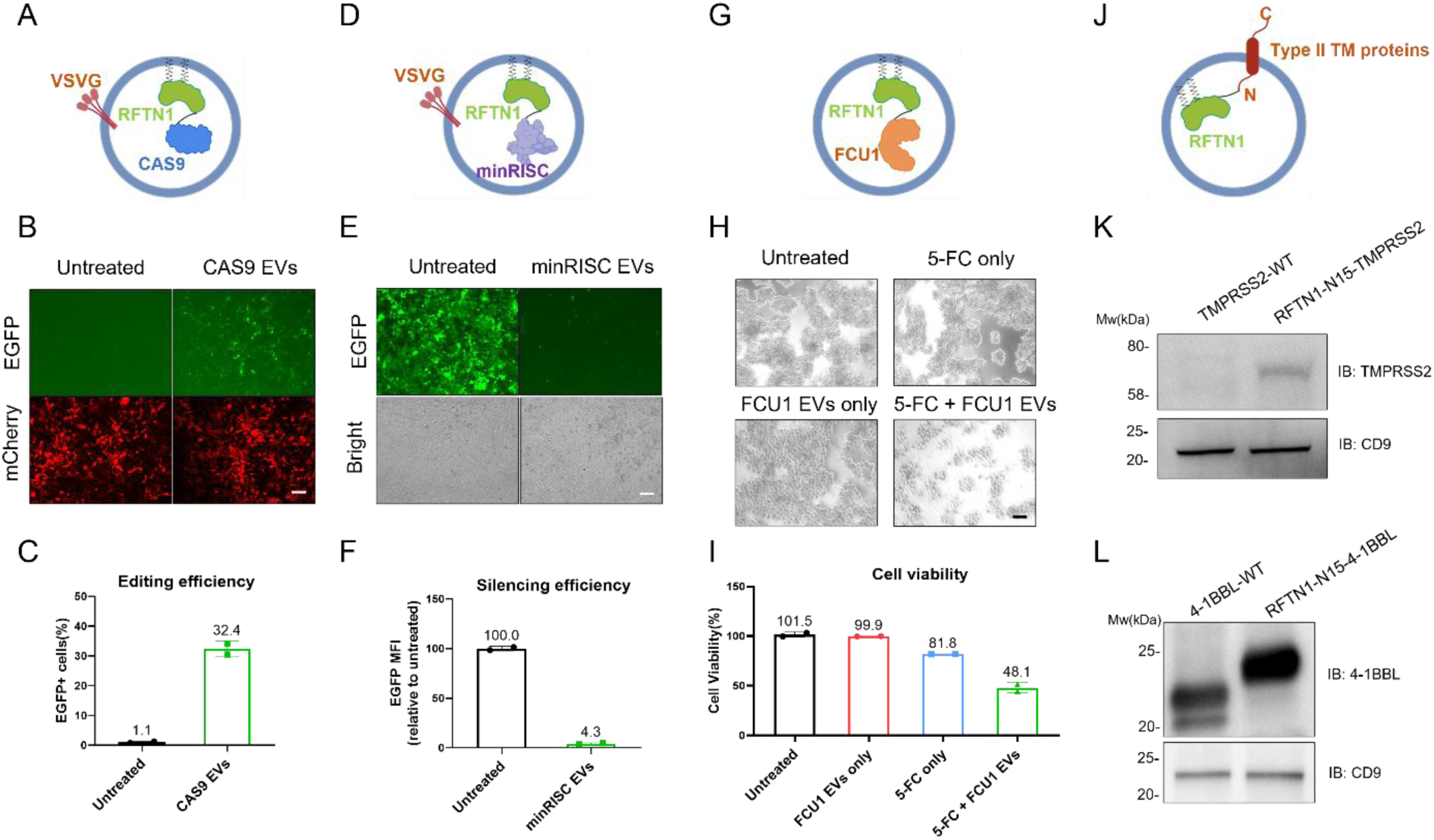
RFTN1-N15 demonstrated versatility in EVs engineering and cargo loading. (A) Schematic diagram showing the EVs engineering of Cas9 mediated by RFTN1-N15. (B) Cas9 reporter assay analysis at 48 hours post EVs treatment. Scale bar: 20 μm. (C) Cas9 editing efficiency analysis as quantified by EGFP positive cells ratio from flow cytometry analysis. N=2. Values are plotted as mean ± SD. (D) Schematic diagram showing the EVs engineering of the minimal RISC comlpex mediated by RFTN1-N15. (E) EGFP silencing analysis at 48 hours post EVs treatment. Scale bar: 20 μm. (F) EGFP silencing efficiency analysis as quantified by EGFP mean fluorescence intensity from flow cytometry analysis. N=2. Values are plotted as mean ± SD. (G) Schematic diagram showing the EVs engineering of FCU1 mediated by RFTN1-N15. (H) Cellular toxicity assay analysis at 48 hours post EVs treatment. Scale bar: 100 μm. (I) Cell viability as quantified by CCK8 assay. N=2. Values are plotted as mean ± SD. (J) Schematic diagram showing the EVs engineering of type II transmembrane proteins mediated by RFTN1-N15. (K) Western blot analysis for engineered TMPRSS2 EVs. (L) Western blot analysis for engineered 4-1BBL EVs.

Next, RFTN1-N15 was further tested for engineering cytoplasm-localized cargos. We have previously designed minRISC-EVs platform as gene silencing tool, which essentially encapsulated EVs with AGO2 protein complexed with guide RNAs (Tao Qiu, 2025). Herein, RFTN1-N15 was fused to the N-terminus of AGO2, while keeping the other elements same as previously described (Figure 3D). More than 95% EGFP silencing rate was observed (total 1E10 particles, 48 hours post treatment), suggesting RFTN1-N15 was capable in loading the gene silencing complex in EVs with high efficiency (Figure 3E-F). In another example, FCU1 was a chimeric protein consisted of yeast cytosine deaminase (CDase) and uracil phosphoribosyltransferase (UPRTase) (Erbs et al., 2000). FCU1 efficiently catalyzed the direct conversion of 5-FC, a relatively nontoxic antifungal agent, into the toxic metabolites 5-fluorouracil (5-FU) and 5-fluorouridine-5’monophosphate (5-FUMP), thus had been investigated for anti-tumor therapy (Erbs et al., 2000). To examine if FCU1 could be loaded into EVs and exhibit function, RFTN1-N15 was fused to N-terminus of FCU1 (Figure 3G). In the presence of 5-FC and engineered EVs, around 50% cell death was observed, confirming the efficacy of the strategy (Figure 3H-I).

Finally, we wondered if RFTN1-N15 could help to enrich type II transmembrane proteins on EVs as well. The transmembrane protease serine 2 (TMPRSS2) was essential host cell factor for aiding the cellular entry of the severe acute respiratory syndrome coronavirus 2 (SARS-CoV-2) (Koyou et al., 2025). Engineered decoy EVs with overexpressed angiotensin-converting enzyme 2 (ACE2) receptor and TMPRSS2 had demonstrated neutralizing effect towards SARS-CoV-2, rendering the decoy EVs to be potential therapeutics (Cocozza et al., 2020). As a type II transmembrane protein, TMPRSS2 had its N-terminus in cytoplasm and EVs lumen side. Therefore, we directly fused RFTN1-N15 to TMPRSS2 N-terminus and analyzed protein level in EVs (Figure 3J). Encouragingly, RFTN1-N15-TMPRSS2 EVs indeed had elevated TMPRSS2 deposition compared to TMPRSSE-WT EVs (Figure 3K). On the other hand, 4-1BBL was a type II transmembrane glycoprotein that served as the ligand for the receptor 4-1BB (Chin et al., 2018; Singh, Kim, Lee, Eom, & Choi, 2024). This interaction played a key role in the immune system by promoting the activation, proliferation, and survival of T cells and NK cells, making it a significant target for immunotherapy (Chin et al., 2018; Singh et al., 2024). EVs engineered to overexpress 4-1BBL had presented prominent anti-tumor effect in cancer therapy (Semionatto et al., 2020). Therefore, RFTN1-N15 was similarly fused to 4-1BBL N-terminus, and the results again confirmed that RFTN1-N15-4-1BBL EVs had increased 4-1BBL density versus 4-1BBL-WT EVs (Figure 3L).

In summary, RFTN1-N15 demonstrated versatility in EVs engineering, allowing efficient cargo loading including nucleus-localized proteins, cytoplasmic proteins as well as type II transmembrane proteins.

### 3.5 Arginase1-loaded EVs demonstrated anti-inflammatory activity both *in vitro* and *in vivo*

In macrophages, the enzyme arginase 1 (ARG1) and inducible nitric oxide synthase (iNOS) competed for the amino acid arginine which determined macrophage function and polarization (Chen et al., 2023). ARG1 broke down arginine into ornithine and urea leading to tissue repair and inflammation inhibition (M2 state), whereas iNOS produced nitric oxide to promote inflammation (M1 state). This polarization balance was crucial for various immune processes, with dysregulation of these enzymes contributing to inflammatory diseases or impaired immunity against pathogens (Chen et al., 2023; Luo, Zhao, Cheng, Su, & Wang, 2024; Orecchioni, Ghosheh, Pramod, & Ley, 2019). We were curious if EVs could be engineered to deliver these two enzymes to macrophages, in order to control the polarization fate of macrophages towards desired directions.

To begin with, RFTN1-N15 was fused to the N-terminus of mouse ARG1 with a flexible GS linker (Figure 4A). Since it was reported that ARG1 assembled as trimer for catalytic activity (Kanyo, Scolnick, Ash, & Christianson, 1996), we reasoned that adding a trimerization motif to the RFTN1-N15-ARG1 fusion protein could potentially help to boost enzymatic activity. Hence, a foldon motif was also added to the C-terminus of the fusion protein (Meier, Guthe, Kiefhaber, & Grzesiek, 2004). Both mARG1 and mARG1-foldon EVs were generated, which presented similar size distribution and morphology (Figure 4A-C). On western blots, it was clear that mARG1-foldon had slightly increased protein weight than the non-foldon version as expected (Figure 4D). To validate the effect of engineered EVs on macrophages, RAW264.7 cells were first polarized towards M1 phenotype by induction with lipopolysaccharide (LPS) and interferon-γ (IFN-γ). Next, EVs were added to those M1 cells, followed by qPCR analysis for gene expression changes at 48 hours post treatment. Interestingly, mARG1-foldon EVs, but not mARG1 EVs, presented dose-dependent suppression effect of pro-inflammatory cytokines including IL-1β and IL-12 (Figure 4E). The observation indicated successful anti-inflammatory modulating effect by ARG1 EVs as expected, and that the trimerization motif was indispensable for the engineering.

**Figure 4.**
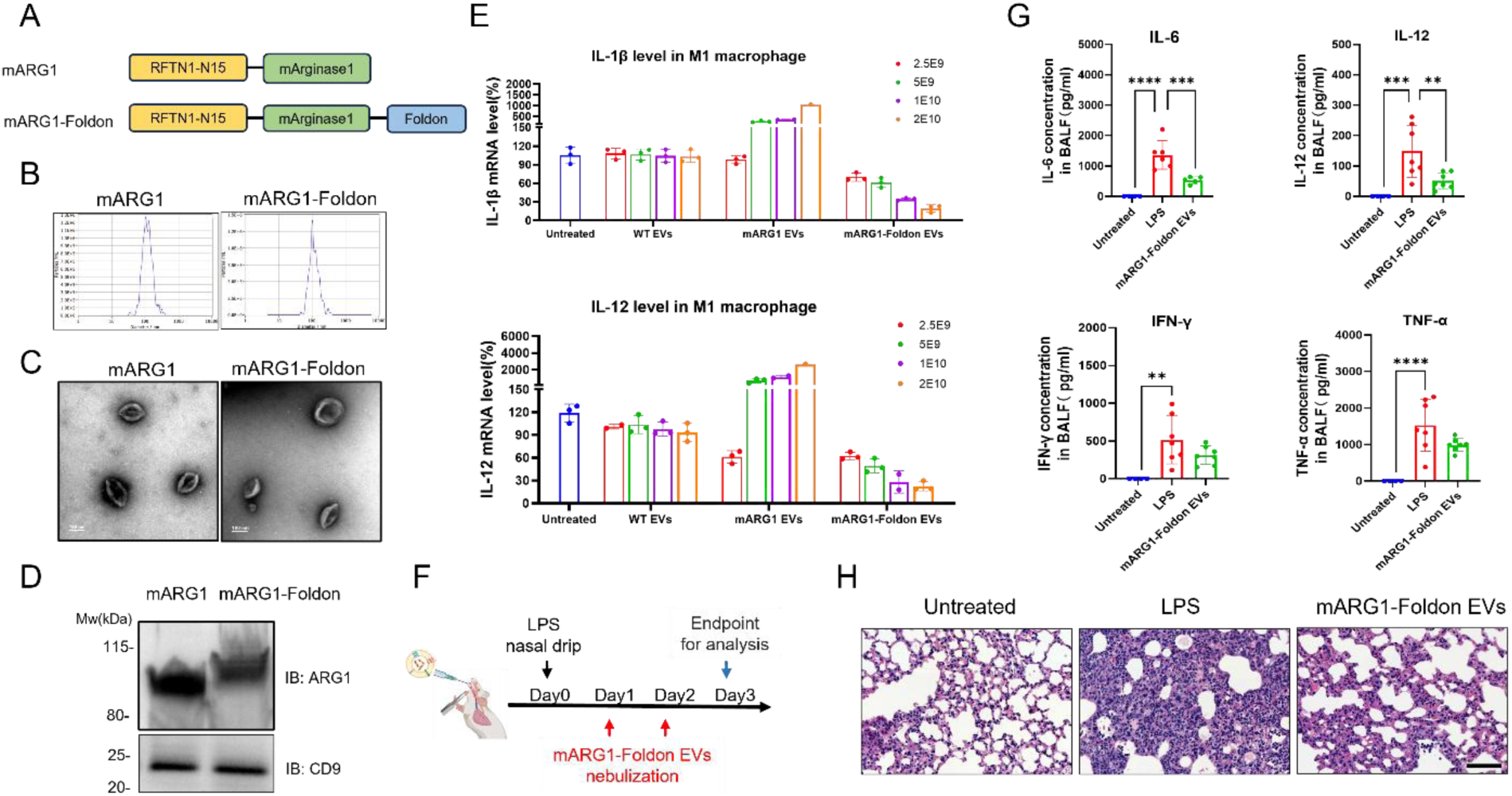
Arginase1-loaded EVs demonstrated anti-inflammatory activity both *in vitro* and *in vivo*. (A) Schematic diagram showing the EVs engineering of mouse arginase1 protein mediated by RFTN1-N15 and foldon trimerization motif. (B) Nanoparticle tracking analysis for purified EVs. (C) Representative transmission electron micrographs of purified EVs. Scale bar: 100 nm. (D) Western blot analysis for purified EVs. (E) QPCR analysis for IL-1β and IL-12 mRNA changes in M1 macrophages receiving different treatment groups. Four doses of EVs from total 2.5E9 particles to 2E10 particles were tested. N=3. Values are plotted as mean ± SD. (F) Schematics of acute lung injury model in C57BL/6J mice. A single dose of LPS nasal drip was administrated at day 0, followed by EVs administration for two consecutive days by nebulization. Mice were euthanized at day 3 for analysis. (G) Pro-inflammatory cytokines analysis in BALF from different groups. N=6. **: p value < 0.01;***: p value < 0.001;****: p value < 0.0001. Values are plotted as mean ± SD. (H) H&E analysis for lung sections from different groups. Scale bar: 100 μm.

To confirm that mARG1-foldon EVs had consistent activity *in vivo*, acute lung injury model was established in C57BL/6J mice by a single dose of LPS nasal drip, followed by EVs treatment for two consecutive days through nebulization (Figure 4F). LPS treatment induced high level of inflammatory cytokines in bronchoalveolar lavage fluid (BALF), whereas mARG1-foldon EVs treatment effectively reduced these cytokines level (Figure 4G). Hematoxylin and eosin (H&E) staining for lung sections revealed that LPS resulted in apparent alveolar edema and massive infiltration of lymphocytes into alveolar regions, whereas mARG1-foldon EVs treatment significantly alleviated these histopathological changes (Figure 4H). Therefore, mARG1-foldon EVs were effective in alleviating acute lung inflammation *in vivo*.

### 3.6 Inducible NOS-loaded EVs demonstrated pro-inflammatory activity both *in vitro* and *in vivo*

On the other hand, to engineer iNOS EVs, RFTN1-N15 was fused to the N-terminus of mouse iNOS with a flexible GS linker (Figure 5A). The catalytically active iNOS was in homodimer form (Ghosh & Stuehr, 1995), hence a dimer version was designed by adding GCN4 dimer motif (Harbury, Zhang, Kim, & Alber, 1993) to the C-terminus of fusion protein (Figure 5A). No obvious size or morphology differences were observed between iNOS-EVs and iNOS-GCN4 EVs, except for slightly increased protein size as expected (Figure 5B-D). To determine EVs function in cell model, RAW264.7 cells were polarized towards M2 phenotype by induction with IL-4, followed by treatment with engineered EVs. Dose-dependent elevation of inflammatory cytokines expression were observed for iNOS-GCN4 EVs treatment, whereas iNOS EVs had low activity (Figure 5E). In conclusion, efficient pro-inflammatory modulation effect by the engineered EVs were as expected, and that the dimerization motif was essential for the engineering.

**Figure 5.**
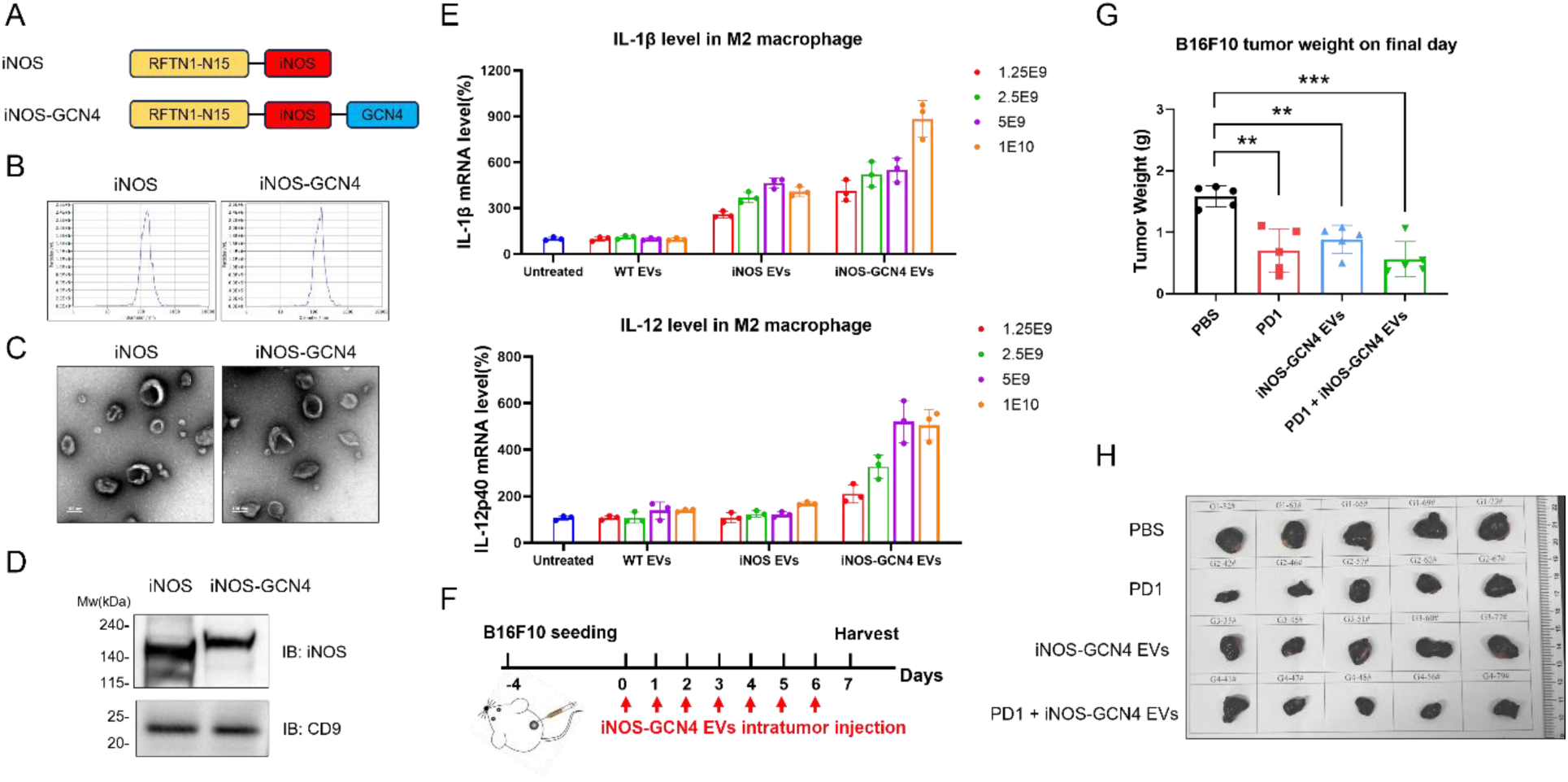
Inducible NOS-loaded EVs demonstrated pro-inflammatory activity both *in vitro* and *in vivo*. (A) Schematic diagram showing the EVs engineering of mouse iNOS protein mediated by RFTN1-N15 and GCN4 dimerization motif. (B) Nanoparticle tracking analysis for purified EVs. (C) Representative transmission electron micrographs of purified EVs. Scale bar: 100 nm. (D) Western blot analysis for purified EVs. (E) QPCR analysis for IL-1β and IL-12 mRNA changes in M2 macrophages receiving different treatment groups. Four doses of EVs from total 1.25E9 particles to 1E10 particles were tested. N=3. Values are plotted as mean ± SD. (F) Schematics of B16F10 xenograft model in C57BL/6J mice. B16F10 cells were first implanted subcutaneously and grew to appropriate size, followed by EVs intratumorally injection for 7 consecutive days. Mice were euthanized at day 7 for analysis. (G) Tumor mass measurement at end point for different treatment groups. N=5. **: p<0.01; ***: p value < 0.001. Values are plotted as mean ± SD. (H) Images of tumor at end point for different treatment groups. N=5.

For validating the consistent efficacy *in vivo*, B16F10 melanoma xenograft model was established in C57BL6/J mice. The iNOS-GCN4 EVs were intratumorally injected for seven days consecutively (Figure 5F). Immune checkpoint inhibitors PD1 antibody was included as monotherapy, as well as in combination therapy with EVs to see if a synergistic effect could be produced. B16F10 tumor expanded rapidly in PBS treated groups, whereas iNOS-GCN4 EVs significantly inhibited tumor growth as measured by tumor size and tumor mass (Figure 5G-H). Notably, iNOS-GCN4 EVs monotherapy resulted in similar tumor inhibition efficacy to PD1 monotherapy, although the combination therapy did not lead to greater inhibitory effect (Figure 5G-H). In summary, iNOS-GCN4 EVs demonstrated anti-tumor efficacy *in vivo*.

### 3.7 Identification of type I transmembrane proteins for EVs surface display

We next moved on to identify scaffold proteins for displaying cargos on EVs surface. Particularly, type I transmembrane proteins as exemplified by PTGFRN and PLXNA1 had been validated for applicability, which also appeared as top enriched candidates in our EVs proteome (data not shown). Structurally, type I transmembrane proteins had signal peptide that directed the secretion of N-terminus to extracellular region, where the EVs surface cargos could be appropriately placed. Therefore, we focused on type I transmembrane proteins as potential candidates. Since the length of PTGFRN (879 aa) and PLXNA1 (1896 aa) were not desirable for the ease of engineering, we set out to screen for candidates that had shorter protein length.

The poliovirus receptor (PVR, 417 aa) was filtered out as potential candidate (Figure 6A). As a type I transmembrane protein, PVR was initially identified as the receptor for the human poliovirus (Mendelsohn, Wimmer, & Racaniello, 1989), and was later found to be essential in mediating cell adhesion, as well as regulating immune response (Bowers, Readler, Sharma, & Excoffon, 2017). Structurally, PVR belonged to the immunoglobulin (Ig) superfamily (Mendelsohn et al., 1989), as it contained several immunoglobulin (Ig)-like domains in tandem at N-terminus (Figure 6A).

**Figure 6.**
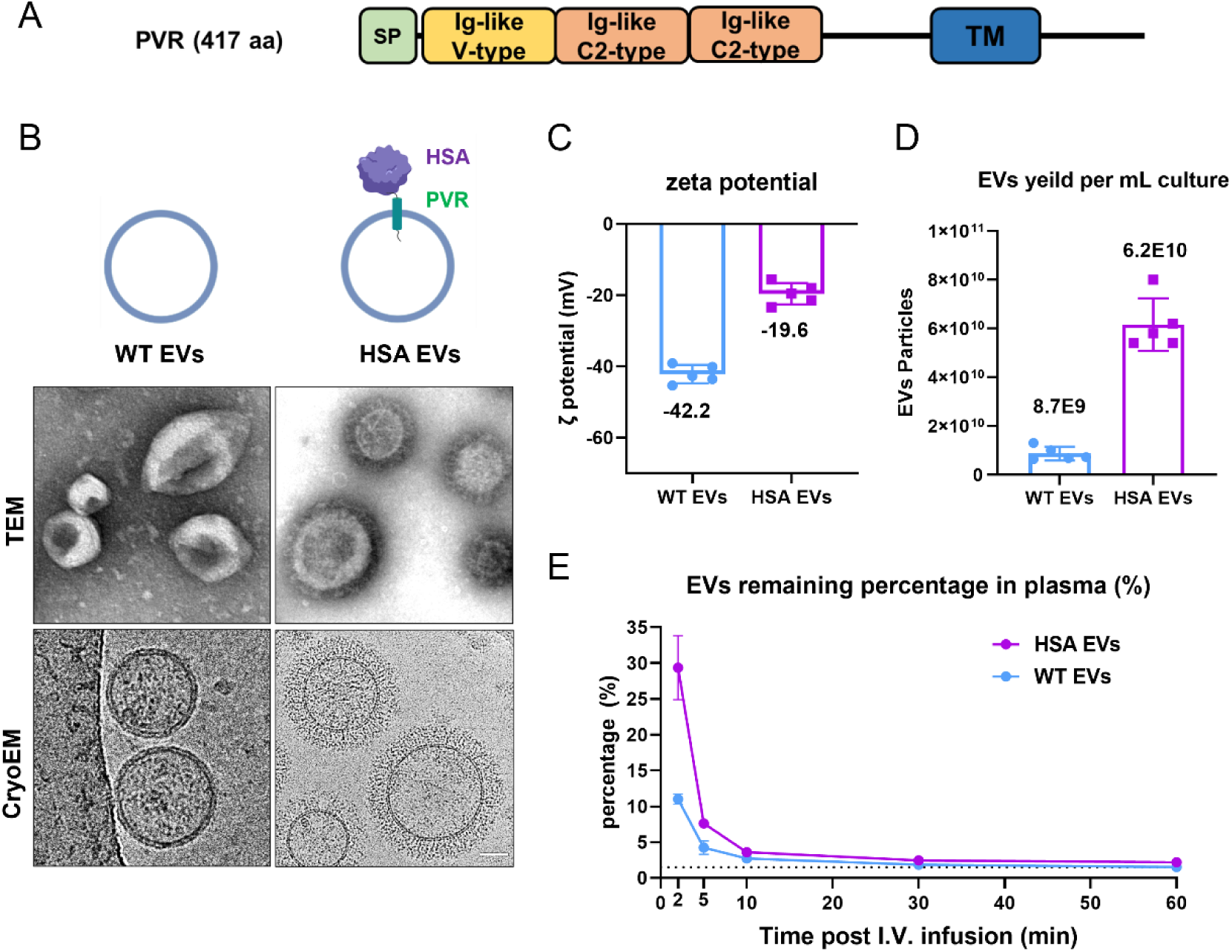
PVR displayed human serum albumin on EVs surface with high density, and extended EVs circulation half-life. (A) Schematic diagram showing the domain organization of PVR. (B) Upper panel: schematic diagram showing the EVs engineering of human serum albumin (HSA) through PVR scaffold. Lower panel: TEM and cryo-EM analysis for EVs. Scale bar: 100 nm. (C) Zeta potential analysis for WT EVs versus HSA EVs. N=5. Values are plotted as mean ± SD. (D) Yield analysis for WT EVs versus HSA EVs. N=5. Values are plotted as mean ± SD. (E) The EVs were intravenously injected in C57BL/6 mice, and the quantity of remaining EVs in the plasma was analyzed at different time points. N=2. Values are plotted as mean ± SD.

### 3.8 PVR displayed human serum albumin on EVs surface with high density, and extended EVs circulation half-life

To analyze PVR’s ability for displaying EVs surface cargos, human serum albumin (HSA) was first selected for validation for following reasons: (1) HSA was secreted protein, thus could be placed in between PVR’s secretion signal and Ig-like domains for EVs surface display; (2) HSA was popular platform for drug half-life extension, due to its interaction with the recycling receptor neonatal Fc receptor (FcRn) (Sleep, Cameron, & Evans, 2013). Upon entering the circulation, EVs were reported with short half-life due to the rapid clearance by liver and spleen (Imai et al., 2015; Lai et al., 2014; Liang et al., 2022; Matsumoto et al., 2020). Therefore, we tested if displaying HSA directly on EVs surface could help to increase EVs retention time in circulation (Figure 6B). A mutant form of human serum albumin (HSA K573P) was selected for higher affinity with FcRn receptor (Andersen et al., 2014). Surprisingly, the display of HSA on EV surface by PVR were observed to be at extremely high-density, as the protein corona-like structures were apparent to coat the EVs surface (Figure 6B). As a result, the zeta-potential of EVs changed significantly, from averaging −40 mV of non-modified EVs, to averaging −20 mV for HSA EVs (Figure 6C). Another interesting observation was that the average yield of EVs also increased around 3- to 5-fold, when HSA was modified on EVs surface (Figure 6D).

The HSA EVs were labeled with DiR fluorescent dye and intravenously injected in C57BL/6 mice. The quantity of remaining EVs in the plasma was analyzed at different time points. As shown in Figure 6E, the amount of HSA EVs in the plasma was average 2- to 3-fold higher to that of WT EVs at early time points (2 min and 5 min), suggesting that HSA modification successfully extended EVs retention time as expected. Nonetheless, beyond 15 min, the remaining EVs in the plasma were still too low to be detected (Figure 6E).

### 3.9 PVR displayed antibodies on EVs surface, and rendered EVs targeting specificity

To further validate the functionality of PVR, we moved on to test additional EVs surface cargos. Displaying antibodies on EVs surface was classic strategy for altering EVs’ targeting ability, and this strategy was analyzed independently in our setting for efficiency. Atezolizumab and trastuzumab were approved antibody drugs which targeted PD-L1 and HER2 receptor, respectively (Shah, Kelly, Liu, Choquette, & Spira, 2018; Stebbing, Copson, & O’Reilly, 2000). Single-chain variable fragments (scFv) of either antibody were constructed and displayed on EVs surface, by direct fusion with PVR scaffold (Figure 7A). The EVs endocytosis analysis was subsequently performed in two cancer cell lines: MDA-MB-231 and SKOV3. SKOV3 expressed both PD-L1 and HER2 receptors on cell surface (Ying et al., 2025), whereas MDA-MB-231 only expressed PD-L1 receptor with high abundancy on surface (Mittendorf et al., 2014). Equal number of DiO dye-labeled EVs particles were added to cells, followed by fluorescent imaging and flow cytometry analysis at two hours post treatment. Compared to non-modified WT EVs, the αPD-L1 EVs had around 10-fold elevation of endocytosis rate in MDA-MB-231 cells, whereas the αHER2 EVs had no increase (Figure 7A-B). In contrast, both αPD-L1 and αHER2 EVs had around 4-fold higher endocytosis rate in SKOV3 cells (Figure 7A-B). The results indicated that PVR successfully displayed antibodies on EVs surface, and importantly rendered EVs with desired targeting ability *in vitro*.

**Figure 7.**
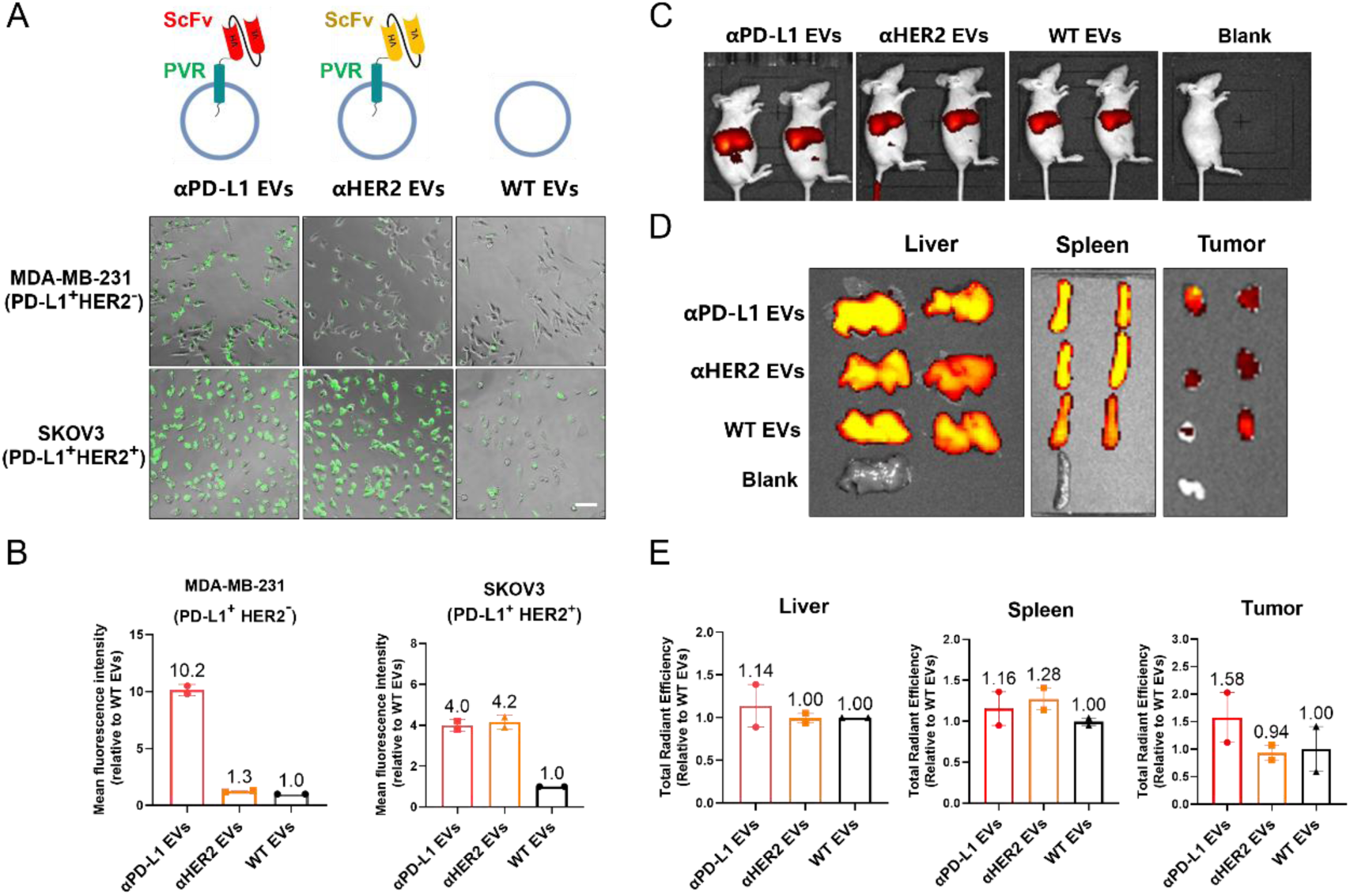
PVR displayed antibodies on EVs surface, and rendered EVs targeting specificity. (A) Upper panel: schematic diagram showing the EVs engineering of single-chain variable fragments (scFv) through PVR scaffold. Lower panel: endocytosis analysis for MDA-MB-231 (PD-L1^+^ HER2^-^) and SKOV3 (PD-L1^+^ HER2^+^) at two hours post EVs treatment. Scale bar: 10 μm. (B) Relative endocytosis efficiency quantified by mean fluorescence intensity from flow cytometry. N=2. Values are plotted as mean ± SD. (C) The nude mice were implanted with MDA-MB-231 xenograft, and imaged at four hours post EVs I.V. infusion. (D) The liver, spleen and tumor tissues were imaged at four hours post EVs I.V. infusion. (E) The amount of EVs in each tissue as quantified by total radiant intensity. Values are plotted as mean ± SD.

To analyzed if the EVs targeting ability was consistent *in vivo*, MDA-MB-231 tumor xenograft model was established in nude mice. The αPD-L1 EVs, αHER2 EVs and WT EVs were labeled with DiR dye, followed by intravenously injection at equal particle number (Figure 7C). At four hours post treatment, the EVs distribution in mice were imaged and quantified by IVIS. Interestingly, the αPD-L1 EVs demonstrated more than 50% higher accumulation in tumors when compared to both αHER2 EVs and WT EVs (Figure 7D-E). Therefore, the antibody-modified EVs retained targeting ability *in vivo* as well.

### 3.10 Comparison of PTGFRN and PVR for EVs cargo loading efficiency

Finally, we compared the performance between PVR and the previously reported scaffold protein PTGFRN. Western blot analysis for HSA-PTGFRN and HSA-PVR EVs revealed that the fusion protein sizes were as expected (Supplementary Figure 2A). ELISA quantification revealed that PVR was able to display around 100 HSA molecules per EV, which was higher than PTGFRN of around 70 (Supplementary Figure 2B). Furthermore, interleukin-15 (IL15) was selected as another cargo which represented secreted cytokines. The fusion proteins sizes presented on WB were also as expected (Supplementary Figure 2C). ELISA quantification for IL15 revealed that PVR was able to display around 4.3 IL15 molecules per EV, which was again higher than PTGFRN of around 1.9 (Supplementary Figure 2D). We concluded that PVR had better performance than PTGFRN in EVs engineering.

## 4. DISCUSSION

EVs as novel therapeutic modality have presented promising therapeutic potential. Engineering EVs for cargo loading and additional gain of functions critically rely on EVs scaffold proteins (Ma et al., 2025; Yang et al., 2024). Several studies have identified novel EVs scaffold proteins based on EVs proteomics, with differences in EVs purification methodology and candidates sorting criteria (Dooley et al., 2021; Zhao et al., 2024; Zheng et al., 2023). Here, we utilized density gradient-based method for EVs purification, and performed differential expression-based proteomics analysis for identifying novel EVs scaffolds being enriched from the rest of cellular proteins. As such, RFTN1 was identified as novel scaffold for EVs luminal loading, and PVR for EVs surface display. A variety of cargoes, including enzymes, gene editing tools, transmembrane proteins, serum albumins and antibodies were successfully engineered onto the surface or into the lumen of EVs at biologically active levels.

Mechanistically, we reported that RFTN1 relied on three signals for efficient EVs sorting ability: myristoylation on glycine 2, palmitoylation on cysteine 3 and a stretch of positively charged residues from lysine 7 to arginine 15. In previous reports, MARCKS relied on a single N-terminus lipidation (Dooley et al., 2021); BASP1 utilized one lipidation anchor and four lysines at N-terminus (Dooley et al., 2021); Src kinase relied on multiple lipid anchors at N-terminus (Whitley et al., 2022). It appeared that RFTN1 had a more balanced combination of amino acid lipidation and charges for membrane association. Accordingly, a minimal sequence derived from the N-terminus 15 amino acids from RFTN1, were found to be sufficient for EVs cargo engineering. We believe RFTN1-N15 is quite versatile for use as EVs engineering tool.

During the EVs engineering process of metabolic enzymes including ARG1 and iNOS, we realized that it was essential to consider protein conformation for retaining enzymatic activity. Although a relatively long and flexible GS linker was used to fuse RFTN1-N15 with ARG1 or iNOS, the fusion protein delivered by EVs was not active unless the appropriate oligomerization motifs were further added. This observation should be instructive for EVs engineering of enzyme cargos in the future.

Type I transmembrane proteins including PTGFRN and PLXNA1 have been reported as EVs scaffolds for displaying surface cargos (Dooley et al., 2021; Zhao et al., 2024), yet both proteins have relatively long length, imposing significant difficulties in construct building and engineering. We reported another type I transmembrane proteins PVR, with significant shorter length while keeping efficient EVs sorting ability. Interestingly, both PVR and PTGFRN belonged to the immunoglobulin (Ig) superfamily (Dooley et al., 2021; Mendelsohn et al., 1989). The observation suggests that proteins from this superfamily may have advantageous structural features for EVs enrichment, which could deserve deeper investigation.

Developing EVs into therapeutics are currently facing several obstacles, with one of these being short circulation half-life (Imai et al., 2015; Lai et al., 2014; Matsumoto et al., 2020). A previous report decorated EVs surface with serum albumin binding domain, such that the EVs would bind serum albumin upon entering the circulation, which effectively extended EVs’ circulation half-life (Liang et al., 2022). In our report, we directly displayed human serum albumins (HSA) on EVs surface, which strategy also prolonged EVs’ circulation half-life. Interestingly, HSA were found to be displayed on EVs surface with high density, and that the serum albumin engineered EVs had higher production yield. Therefore, serum albumin-displayed EVs could potentially serve as a modular platform with higher stability and lower production cost, which ultimately may benefit the translation of EVs into therapeutics.

## ACKNOWLEDGMENTS

Some illustrative graphics were created with biogdp.com. This study was funded by Vesicure Therapeutic.

## AUTHORSHIP CONTRIBUTION STATEMENT

Tao Qiu: Writing–original draft, Investigation, Data curation, Conceptualization. Rui Hu: Investigation. Yuan Yi: Investigation. Wenqiang Lu: Investigation. Chuang Cui: Investigation. Shuiqin Niu: Investigation. Ke Xu: Writing–review & editing, Supervision.

## DECLARATION OF COMPETING INTEREST

The authors declare that they have no known competing financial interests or personal relationships that could have appeared to influence the work reported in this paper.

---

**Supplementary Figure 1.**
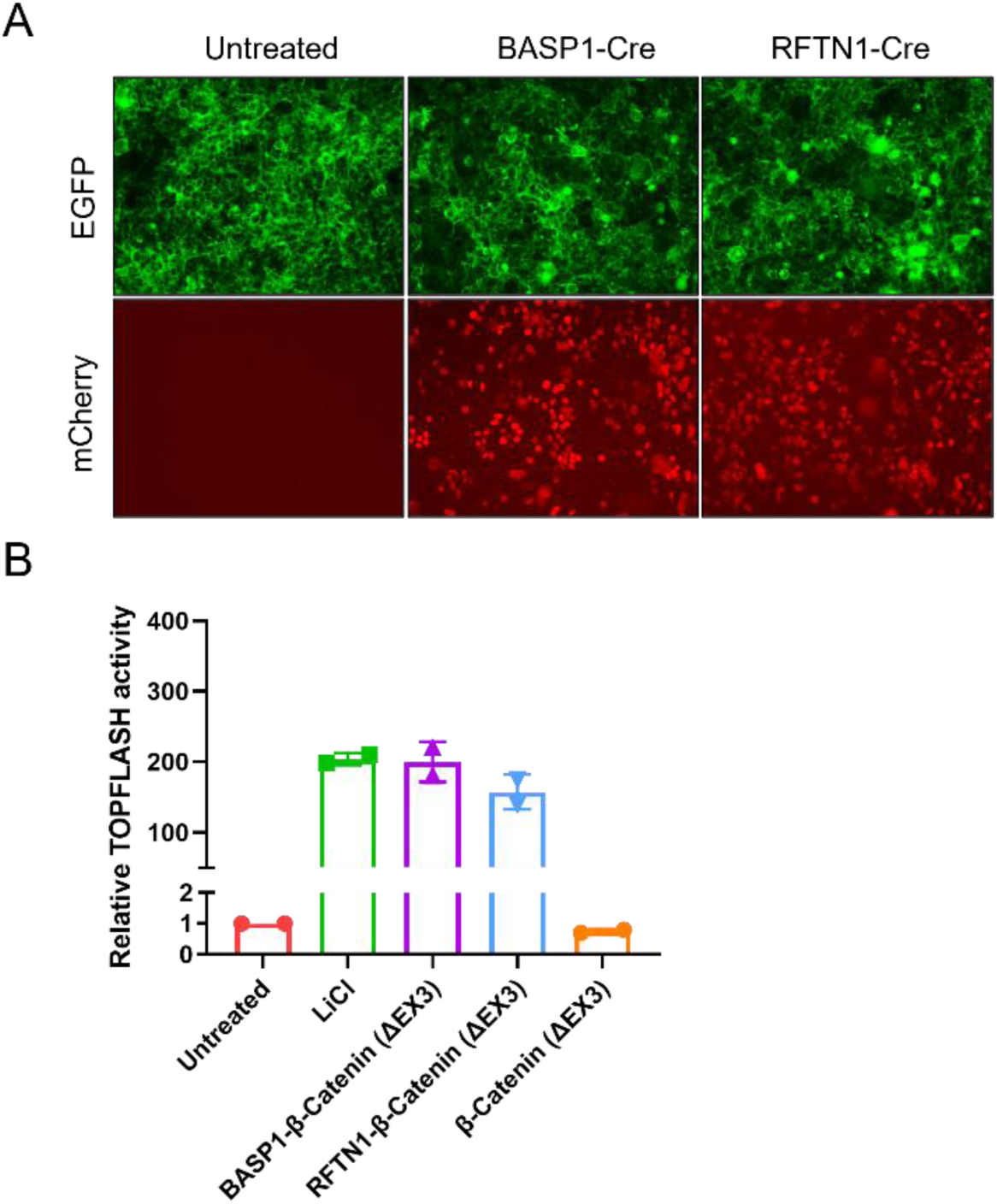
Comparison of BASP1 and RFTN1 in EVs engineering efficiency. (A) Cre reporter assay analysis at 48 hours post EVs treatment. BASP1-Cre and RFTN1-Cre EVs were added to reporter cells at equal quantity (1E10 EVs particles were added to 3E4 reporter cells). (B) TOPFLASH reporter assay analysis at 48 hours post EVs treatment. BASP1-β-catenin (ΔEX3) and RFTN1-β-catenin (ΔEX3) EVs were added to reporter cells at equal quantity (1E10 EVs particles were added to 3E4 reporter cells). N=2. Values are plotted as mean ± SD.

**Supplementary Figure 2.**
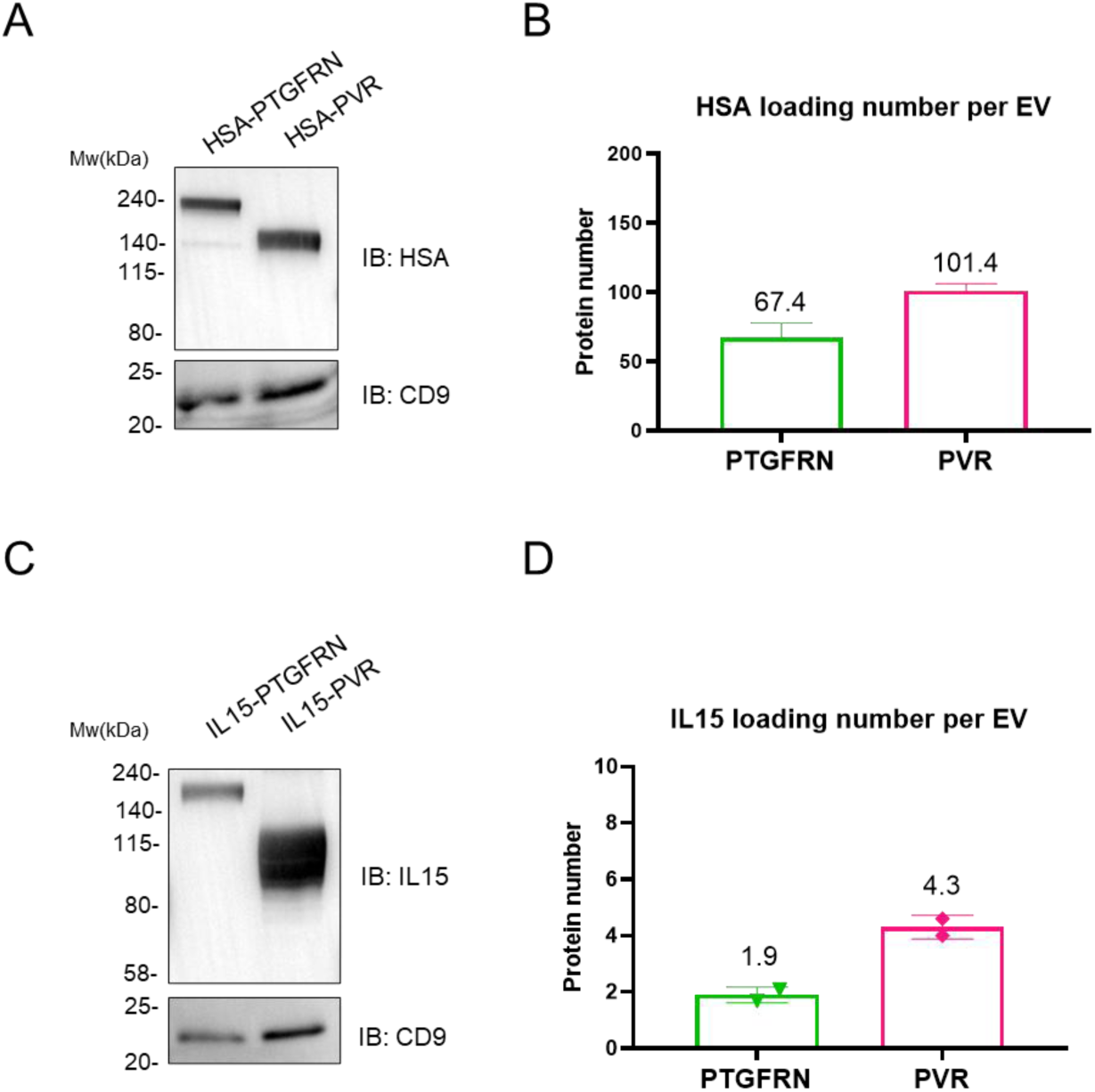
Comparison of PTGFRN and PVR in EVs engineering efficiency. (A) Western blot analysis for purified EVs loaded with HSA by PTGFRN and PVR. (B) ELISA quantification analysis for HSA loading number per EV. N=2. Values are plotted as mean ± SD. (C) Western blot analysis for purified EVs loaded with IL15 by PTGFRN and PVR. (D) ELISA quantification analysis for IL15 loading number per EV. N=2. Values are plotted as mean ± SD.

## Notes

### Competing Interest Statement

The authors have declared no competing interest.

